# Susceptibility of tree shrew to SARS-CoV-2 infection

**DOI:** 10.1101/2020.04.30.029736

**Authors:** Yuan Zhao, Junbin Wang, Dexuan Kuang, Jingwen Xu, Mengli Yang, Chunxia Ma, Siwen Zhao, Jingmei Li, Haiting Long, Kaiyun Ding, Jiahong Gao, Jiansheng Liu, Haixuan Wang, Haiyan Li, Yun Yang, Wenhai Yu, Jing Yang, Yinqiu Zheng, Daoju Wu, Shuaiyao Lu, Hongqi Liu, Xiaozhong Peng

## Abstract

Since SARS-CoV-2 became a pandemic event in the world, it has not only caused huge economic losses, but also a serious threat to global public health. Many scientific questions about SARS-CoV-2 and COVID-19 were raised and urgently need to be answered, including the susceptibility of animals to SARS-CoV-2 infection. Here we tested whether tree shrew, an emerging experimental animal domesticated from wild animal, is susceptible to SARS-CoV-2 infection. No clinical signs were observed in SARS-CoV-2 inoculated tree shrews during this experiment except the increasing body temperature (above 39° C) particular in female animals during infection. Low levels of virus shedding and replication in tissues occurred in all three age groups, each of which showed his own characteristics. Histopathological examine revealed that pulmonary abnormalities were mild but the main changes although slight lesions were also observed in other tissues. In summary, tree shrew is not susceptible to SARS-CoV-2 infection and may not be a suitable animal for COVID-19 related researches.

---

The manifestation caused by SARS-CoV-2 infection is known as COVID-19. Since the first case of SARS-CoV-2 infection reported in Wuhan, China, it has been about five months. It causes a pandemic with more than three million confirmed cases and nearly 230 thousand deaths in addition to huge economic losses to the world^1^. Nevertheless, it is widely considered to be controllable according to the experiences from China and other countries. However, there are still some critical aspects that need to be further investigated in COVID-19 patients, such as cytokine storm, immunopathogenic damages, tropism of SARS-CoV-2, and other sources of SARS-CoV-2 infection besides bat and pangolin that are regarded as the origin of SARS-CoV-2^2–4^. From these points of view, studies of animals become essentially important. In fact, several animal models of COVID-19 have been recently reported in murine^5^, hamster^6^, ferret^7^, and non-human primate^8–11^, which recapitulate COVID-19 from different aspects. In terms of susceptibility to SARS-CoV-2, in addition to these experimental animals, domestic animals and pets are also investigated^12^. Cats, as a popular pet, could be an important source of SARS-CoV-2 infection due to their close relationship with human beings.

The tree shrew, also known as *Tupaia belangeris*, is genetically demonstrated to be close to primates^13–15^. Therefore, it is being developed to be an experimental animal that could be an alternative to primates in biomedical research due to its unique characteristics^16^. In fact, tree shrew has been used for several animal models of virus infections, including hepatitis B^17^, influenza virus^18–20^, and Zika virus^21^. However, *Tupaia* model of high pathogenic viruses has not been reported yet, including SARS-CoV-2. Several reports show that SARS-CoV-2 may originate from wild animals^2–4^. Replication of SARS-CoV-2 in tree shrews is still unknown. In this study, in order to determine the possibility of tree shrew as a COVID-19 model, we tested the susceptibility of tree shrew to SARS-CoV-2 infection. We found that SARS-CoV-2 had limited replication and shedding in tree shrew, and cause mild histopathology, but no typical symptom is observed in infected tree shrew as reported in COVID-19 patients.

## METHODS

### Experimental animals and Ethics

Tree shew, *Tupaia belangeri chinensis*, were bred in the institute of medical biology, Chinese Academy of Medical Sciences. After quarantine and before infection, animals were transferred to ABSL-3 facility and housed in the isolated ventilation cages, with 12-hour light and 12-hour dark. All animal procedures were approved by the Institutional Animal Care and Use Committee of Institute of Medical Biology, Chinese Academy of Medical Science (Ethics number: DWSP202002 001), and performed in the ABSL-3 facility of National Kunming High-level Biosafety Primate Research Center, Yunnan China.

### Study procedures

Procedures in this study are outlined in Figure 1. Total 24 tree shrews were used in this study, and divided into three groups with consideration of gender and age, including young (6 months to 12 months), adult (2 years to 4 years) and old (5 years to 7 years) groups. Each group contains half male and half female animals.

**Figure 1.**
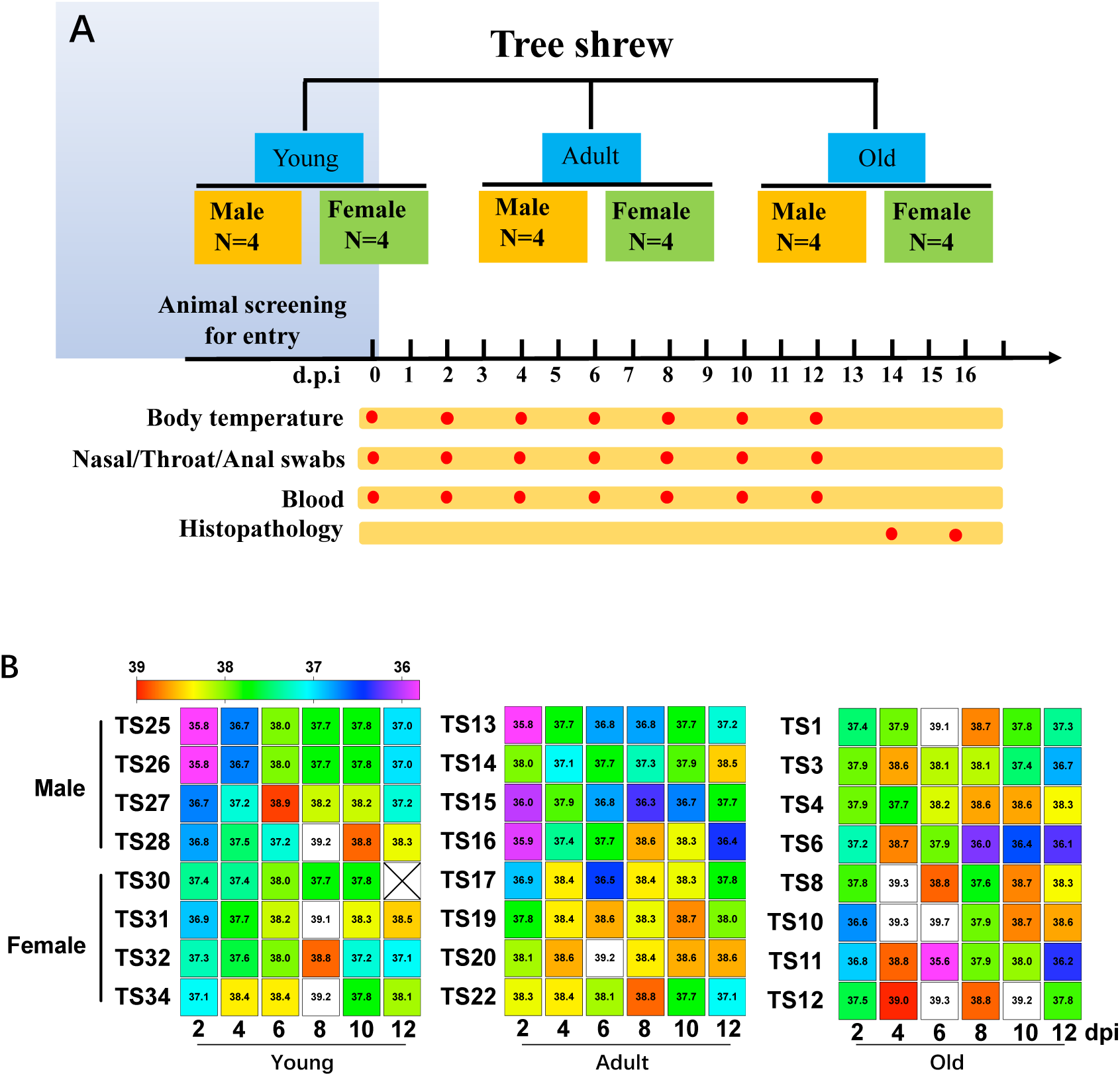
Study design and body temperature in SARS-CoV-2 infected tree shrews. **(A)** Total 24 tree shrews (*Tupaia belangeris*) were divided into three groups (young, adult and old) according to ages. Each group included half male and half female. Each of tree shrew was inoculated with 1ml 10^6^ pfu SARS-CoV-2 nasally (500ul/each) in 0 dpi. Every other day body temperature was monitored. At the same time, nasal, throat, anal and blood samples were collected for analysis of viral loads. On 14 and 16 dpi, animals were euthanatized and necropsied. Gross lesions were recorded and tissue samples were collected for further analysis. **(B)** On every other day as indicated in (A), body temperature of tree shrew was monitored and recorded. The software Graphpad was utilized for data processing and plotting as rainbow heat map. X represents no data collected. Body temperature beyond 39 was shown in white boxes.

After baseline data and samples were collected right before virus inoculation, each of tree shrew was inoculated with 1 ml 10^6^ pfu SARS-CoV-2 nasally (500ul/each nostril). Clinical signs were recorded daily, including behavior, drinking and eating, breathing, feces and so on. Body temperature was also monitored every other day post viral inoculation. At the same time points, nasal, throat and anal swab, and blood samples were collected. Viral genomic RNA in these samples was quantified by RT-qPCR using virus-specific primers and probes. On day 14 post viral inoculation, six animals were anesthetized, bled and necropsied. After gross lesions were recorded, tissue samples were harvested for analysis of viral loads and histopathology.

### Quantification of viral genomic RNA

Swab samples soaked in Trizol solution were vortexed and swabs were removed. 200 µl of each sample were used for RNA extraction and purification via the kit Direct-zol™ RNA MiniPrep. Tissue samples were homogenized in Trizol and 200 µl of each for total RNA preparation. Briefly, 50µl RNase/DNase-free H2O was used to elute RNA from the column. 7.5 µl of each RNA was analyzed in each well of 384-well plate by one-step RT-qPCR using gene-specific primers and probe as described before^8^.

### Histopathological analysis

Tissue samples of heart, liver, spleen, lung, kidney, weasand, stomach, small intestine, rectum, pancreas, brain, spinal cord, uterus, penis, testis, cecum were harvested and fixed in 10% neutral buffered formalin. Paraffin-embedded tissues were cut into 5 µm of sections, followed by haematoxylin and eosin (H&E) staining. Slides were scanned with 3DHISTECH and inspected by the experienced pathologist using the manufacture provided software CaseViewer.

## RESULTS

### Clinical signs in SARS-CoV-2 infected tree shrew

During the period of this study, we couldn’t observe any other clinical sign besides change of body temperature. After SARS-COV-2 inoculation, body temperature of tree shrew was monitored every other day. Among three ages of virus-inoculated tree shrews, all young tree shrews except TS30 showed increasing body temperature. There are 5 young tree shrews (2 males, 3 females) with peak body temperature on 6 or 8-day post inoculation (dpi), followed by gradual decline. The peaks of body temperatures in two female (TS31, TS34) and one male (TS28) young tree shrews were above 39°C (Figure 1A). In addition, among old tree shrews, one male (TS1) and three female (TS8, TS10 and TS12) had the peak body temperature (>39°C) on 4 or 6 dpi. However, only one adult female (TS20) showed the peak body temperature (39.2 °C) on 6 dpi (Figure 1B & C). These results indicated that young/old tree shrews and female tree shrews showed higher sensitivity to SARS-CoV-2 infection, as compared to adult and male tree shrews.

### Virus shedding from SARS-CoV-2 infected tree shrew

On 6 dpi, we could detect genomic RNA of SARS-CoV-2 in 5 nasal swabs, 3 throat swabs, 2 anal swabs and one serum sample from young tree shrews, in 2 nasal swabs from adult tree shrews and in 2 nasal swabs, 1 anal swab from old tree shrew (Table 1). Notably, four samples (nasal, throat, anal swabs and blood) collected from young tree shrew TS27 all showed virus RNA positive. The highest copy number of viral genomic RNA was 10^5.92^/ml in nasal swab from the young tree shrew TS27. On 8 dpi, there were four young, three adult and four old tree shrew with viral RNA positive in some of samples. The highest level of virus shedding was still from the young tree shrew (TS28). On 12 dpi, the young tree shrews showed the decreasing virus shedding and only the animal TS27 had RNA-positive throat swab. In contrast, increasing number of old tree shrews had detectable viral RNA in nasal, throat, and anal swabs (Table 1). From these findings, we deduced that young tree shrew was more susceptible to SARS-CoV-2 infection than adult and old tree shrew. However, old tree shrew and adult tree shrews gave longer duration of virus shedding than young tree shrews. Moreover, we noticed that more male animals of adult and old tree shrews showed virus shedding than females.

**Table 1.**
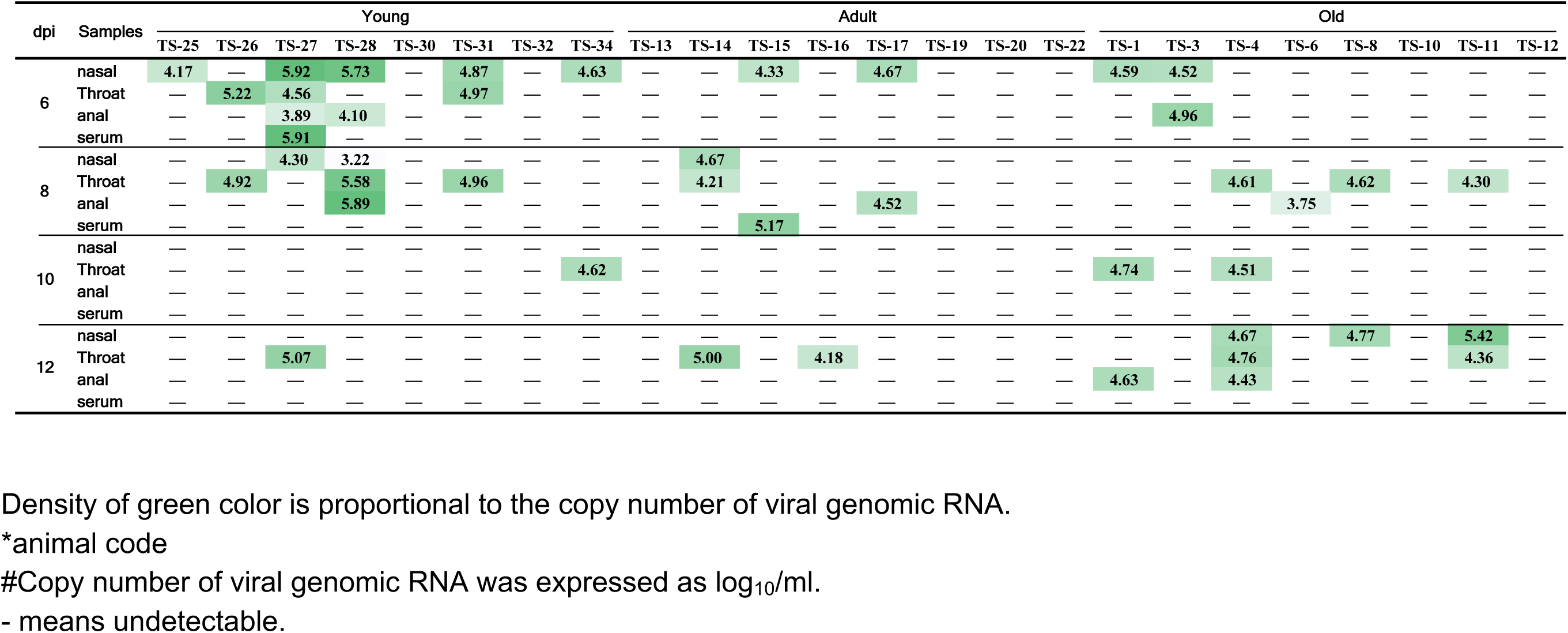
Copy number of viral genomic RNA in clinical samples from SARS-CoV-2 infected tree shrews

### Viral load in tissue samples from SARS-CoV-2 infected tree shrew

In order to determine viral load in tissue samples, we necropsied 8 animals from 3 ages of tree shrews on 14 dpi. Sixteen major tissues were collected from each animal. Viral genomic RNA was quantified as described in Methods. In three young tree shrews (TS26, TS27 and TS28), we could detect viral RNA from only lungs in TS26 and TS27, but not in any tissue from TS28, although these animal had higher number of viral genomic copy numbers at the earlier stage of SARS-CoV-2 infection. In contrast, in the adult tree shrew TS16, six tissues were RNA positive with the highest number 10^9.08^/ml in pancreas. The female adult tree shrew TS14 had viral RNA-positive uterus in addition to lung and pancreas. The other four necropsied tree shrews showed viral RNA-negative results in all 16 tissues (Table S1).

### Histopathological characteristics in SARS-CoV-2 infected tree shrew

To determine the host response to SARS-CoV-2 infection, we examined sixteen tissues from each of 24 tree shrews necropsied on 14 dpi and 16 dpi. No gross lesion was observed in any organ of infected tree shrews. Histopathological inspection revealed that 11 out of 16 tissues had various degrees of pathological changes. The lung of tree shrews in all age groups showed widened pulmonary septum, hyperemia of interstitium, airway obstruction, consolidation of lung margin, local hemorrhagic necrosis, and infiltration of inflammatory cells. Inflammatory cells infiltrated into the submucosa of the trachea in the aged group. Some spleen corpuscles are in active state, and the structure of spleen corpuscles is disordered. Hyperplasia of intestinal gland was observed in the small intestine. The other tissues had certain inflammatory infiltration and massive lymphocyte aggregation (Figure 2). Overall evaluation of histopathology in tissues from all 24 tree shrews revealed that all tissues, except lung, showed only mild histological abnormality. Lung was the major organs affected by SARS-CoV-2 infection. Mild pulmonary abnormality was observed in 62.5%, intermediate abnormality in 12.5% and severe in 4.2% of 24 tree shrews. Unexpectedly, we also found some mild histopathological changes in brain, heart, liver and pancreas (Table 2).

**Figure 2.**
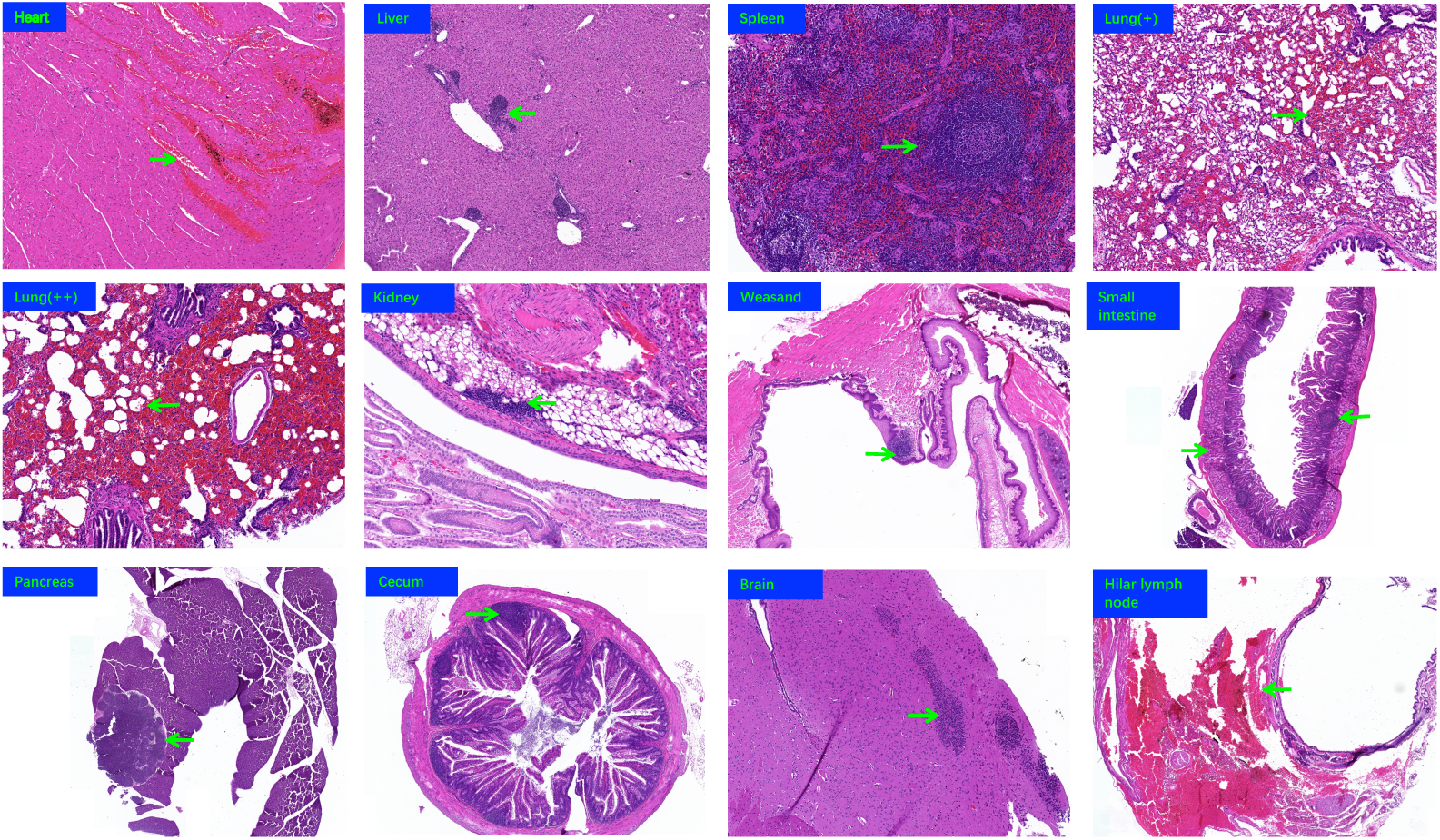
Histopathological examination of affected tissues from SARS-CoV-infected tree shrews. On 14 and 16 dpi, tree shrews were euthanized and necropsied. Seventeen tissues were collected from each animal for histopathological analysis. From eight old tree shrews, eleven tissues with various degrade of pathological changes were representatively shown. Green arrows indicate the pathological changes described in text. Histopathology of all other tree shrews was summarized in Table 2.

**Table 2.**
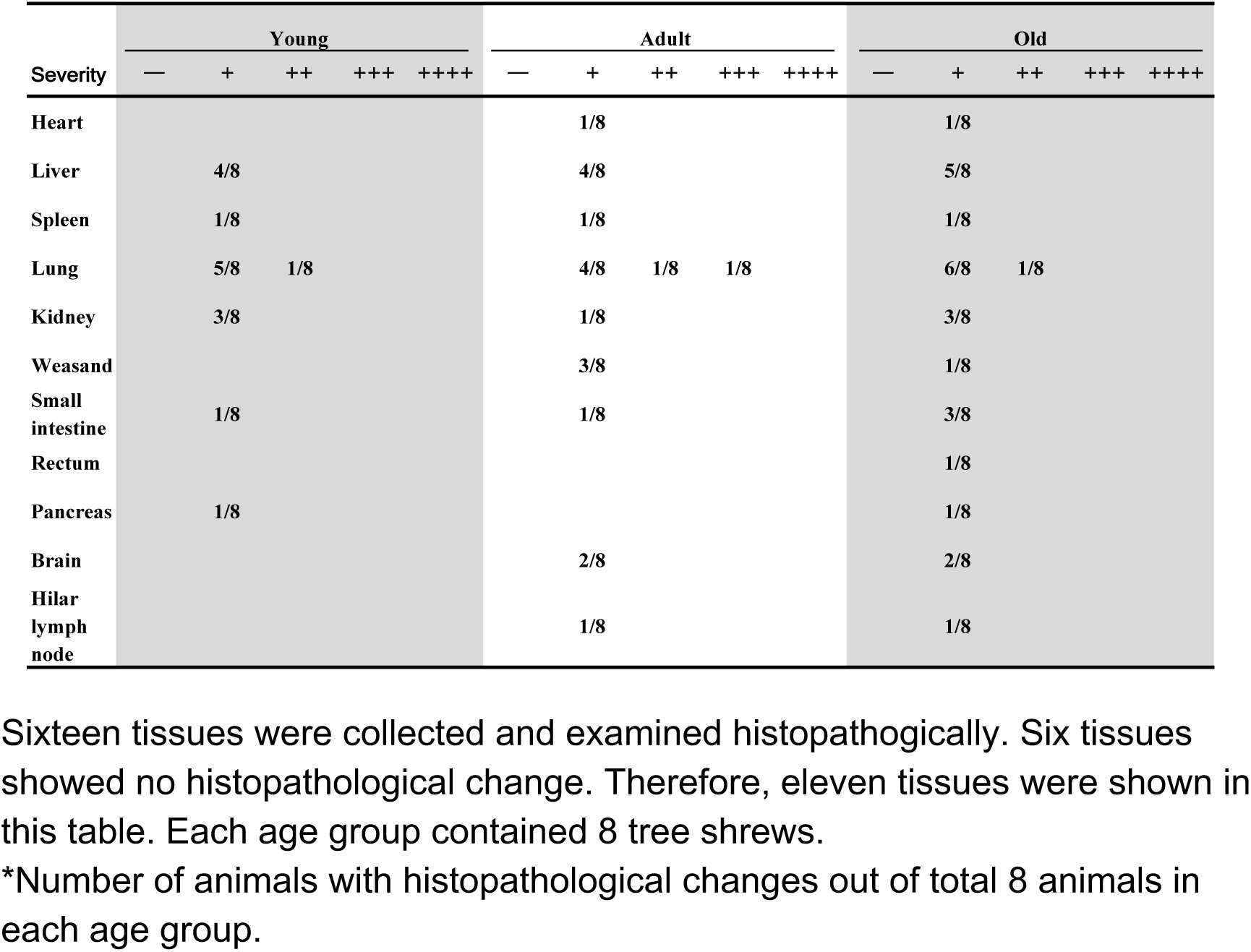
Summary of histopathological examination of tissues from all 24 tree shrews of each age group.

## DISCUSSIONS

The announcement of the genome of SARS-CoV-2 reveals the unique relationship of SARS-CoV-2 with 2003 SARS-CoV and 2012 MERS-CoV^22^, which not only attracts great attentions but also leads to many conjectures. The main concerns are the origin of SARS-CoV-2, the natural reservoir, intermediate host, factors affecting infection and prognosis, and so on. These are actually about the sensitivity or susceptibility of host to SARS-CoV-2 infection^12^. Although the retrospective studies of COVID-19 have given us some answers^23^, further investigation also needs to be performed in various animals. So far, several animal models of COVID-19 have been reported to successfully recapitulate some aspects of this disease^5–11^ and also indicate that these experimental animals are susceptible to SARS-CoV-2 infection. *Rhinolophus sinicus* is mostly recognized as the natural host for SARS-CoV-2^24^. However, the intermediate host is still controversial. Nevertheless, some domestic animals and pets, including cat, can be infected by SARS-CoV-2^12^. Our previous study revealed the variable susceptibility among three species of non-human primates to SARS-CoV-2 infection^8^. Here we experimentally infected tree shrew, an emerging experimental animal genetically close to NHP^14^. No typical clinical sign except the increasing body temperature was observed in SARS-CoV-2 infected tree shrews, although viruses were detectable in swabs, serum samples and some of tissues. These results indicate that tree shrew may be a potential intermediate host for SARS-CoV-2. It is clinically reported that old COVID-19 patients show high morbidity and mortality, which is thought to be associated with the comorbidities^25^. Children confirmed as SARS-CoV-2 infection showed milder clinical symptoms than adult COVID-19 patients ^26^. However, COVID-19 children are reported to be critical in fecal-oral transmission due to persistence of fecal virus shedding even though nasopharyngeal swabs are tested virus-negative ^27^. Results in this study indicate that age may affect SARS-CoV-2 infection of tree shrew. Firstly, more young and old tree shrews showed increasing body temperature than adult tree shrews (Figure 1). Secondly, young tree shrews had more severe virus shedding at the early stage of virus infection than the other animals, and old tree shrews had a longer duration of virus shedding than others (Table 1). One of young male tree shrew, TS27, didn’t have significant increasing body temperature, but virus shedding from nasal, throat, anal and serum was detected on 6 dpi. These results indicated the asymptomatic infection of SARS-CoV-2 in young tree shrews.

Although SARS-CoV-2 infection didn’t cause severe disease in all three ages of tree shrews, viral replication and mild histopathological changes were still observed in this study. Particularly, we found the severe lung abnormality of histology in one adult tree shrew, which may be caused by immunopathological responses, such as cytokine storm ^28^. Unfortunately, reagents for analysis of cytokines in tree shrews are very limited so that we couldn’t further evaluate levels of inflammatory cytokines although we may analyze mRNA levels of cytokines via RT-qPCR in the future. Angiotensin-converting enzyme 2 (ACE2) has been demonstrated as an essential receptor for SARS-CoV-2 infection in vertebrate animals^29^. The interaction between ACE2 and RBD domain of viral spike protein determines the susceptibility and range of host to SARS-CoV-2 infection^30^. Furthermore, distribution of ACE2 *in vivo* may be associated with viral pathogenesis^31^. However, no profile of ACE2 in various tissues of animal models has been reported so far. Amino acid alignment of critical domain of ACE2 binding spike protein RBD showed that there were 10 amino acid different between tree shrew and human, whereas no difference exists between rhesus monkey and human^8^. This may be one of the reasons that tree shrews are not susceptible to SARS-CoV-2 infection.

SARS-CoV-2 distribution of tissues in tree shrew is different from that reported in autopsied COVID-19 patients. For example, high copy number of viral genomic RNA was unexpectedly detected in kidney, pancreas, and spinal cord in SARS-CoV-2 infected tree shrew (Table 3). These results should further be confirmed via nucleic acid *in situ* hybridization, immunohistochemistry or immunofluorescent staining.

In conclusion, tree shrew is not as susceptible to SARS-CoV-2 infection as the reported animal models of COVID-19, though limited replication of SARS-CoV-2 and mild histopathology was detected and observed in some tissues. In addition, commercial reagents and completely domesticated tree shrews are very limited. Therefore, tree shrew is not a suitable experimental animal for COVID-19 related studies. However, it should be very important to investigate whether wild tree shrews in nature are infected by or asymptomatic carrier of SARS-CoV-2.

## Contributors

Shuaiyao Lu, Yuan Zhao, Junbin Wang, Dexuan Kuang, Wenhai Yu, Yun Yang, Haixuan Wang, Haiyan Li and Daoju Wu performed animal-related experiments. Jiahong Gao, Kaiyun Ding and Haiting Long did histopathological work. Jingwen Xu, Mengli Yang, Chunxia Ma, Jingwen Xu, Siwen Zhao, Jingmei Li detected clinical samples. Jing Yang, Yun Yang and Yinqiu Zheng collected and analyzed data. Xiaozhong Peng. Hongqi Liu, Shuaiyao Lu, Yuan Zhao designed the experiments, interpreted data and wrote the manuscript.

## Acknowledgements

The authors would like to thank all staffs in National Kunming High-level Biosafety Primate Research Center for providing ABSL3-related services. This study was supported by 2020YFC0841100 and 2020YFC0846400.

---

**Table S1.** Viral load in tissues collected from SARS-CoV-2 infected tree shrews.

|  | Young |  |  | Adult |  |  |  | Old |
| --- | --- | --- | --- | --- | --- | --- | --- | --- |
|  | TS-26 | TS-27 | TS-28 | TS-13 | TS-14 | TS-15 | TS-16 | TS-12 |
| Heart | - | - | - | - | - | - | - | - |
| Liver | - | - | - | - | - | - | 5.08 | - |
| Spleen | - | - | - | - | - | - | - | - |
| Lung | 5.95 | 5.75 | - | - | 5.04 | - | 6.04 | - |
| Kidney | - | - | - | - | - | - | 8.08 | - |
| Weasand | - | - | - | - | - | - | 8.20 | - |
| Stomach | - | - | - | - | - | - | - | - |
| Small intestine | - | - | - | - | - | - | - | - |
| Rectum | - | - | - | - | - | - | - | - |
| Pancreas | - | - | - | - | 6.67 | - | 9.08 | - |
| Brain | - | - | - | - | - | - | - | - |
| Spinal cord | - | - | - | - | - | - | 4.73 | - |
| Uterus | - | - | - | - | 4.83 | - | - | - |
| Penis | - | - | - | - | - | - | - | - |
| Testis | - | - | - | - | - | - | - | - |
| Cecum | - | - | - | - | - | - | - | - |

Density of green color is proportional to the copy number of viral genomic RNA.
\*animal code

## REFERENCES

1. Coronavirus (COVID-19). https://covid19.who.int/.

2. Susanna, K.P.L., et al. Possible Bat Origin of Severe Acute Respiratory Syndrome Coronavirus 2. Emerging Infectious Disease journal 26(2020).

3. Zhang, T., Wu, Q. & Zhang, Z. Probable Pangolin Origin of SARS-CoV-2 Associated with the COVID-19 Outbreak. Current Biology 30, 1346–1351.e1342 (2020).

4. Lam, T.T.-Y., et al. Identifying SARS-CoV-2 related coronaviruses in Malayan pangolins. Nature (2020).

5. Bao, L., et al. The Pathogenicity of SARS-CoV-2 in hACE2 Transgenic Mice. bioRxiv, 2020.2002.2007.939389 (2020).

6. Chan, J.F.-W., et al. Simulation of the clinical and pathological manifestations of Coronavirus Disease 2019 (COVID-19) in golden Syrian hamster model: implications for disease pathogenesis and transmissibility. Clin Infect Dis 2020 ciaa325, https://doi.org/10.1093/cid/ciaa325.

7. Kim, Y.-I., et al. Infection and Rapid Transmission of SARS-CoV-2 in Ferrets. Cell host & microbe S1931-3128, 30187–30186 (2020).

8. Lu, S., et al. Comparison of SARS-CoV-2 infections among 3 species of non-human primates. bioRxiv, 2020.2004.2008.031807 (2020).

9. Chao, S., et al. Infection with Novel Coronavirus (SARS-CoV-2) Causes Pneumonia in the Rhesus Macaques. research square 27 February 2020, PREPRINT (Version 1) available at Research Square https://doi.org/10.21203/rs.2.25200/v1.

10. Munster, V.J., et al. Respiratory disease and virus shedding in rhesus macaques inoculated with SARS-CoV-2. bioRxiv, 2020.2003.2021.001628 (2020).

11. Rockx, B., et al. Comparative Pathogenesis Of COVID-19, MERS And SARS In A Non-Human Primate Model. bioRxiv, 2020.2003.2017.995639 (2020).

12. Shi, J., et al. Susceptibility of ferrets, cats, dogs, and other domesticated animals to SARS-coronavirus 2. Science, eabb7015 DOI: 7010.1126/science.abb7015 (2020).

13. Fan, Y., et al. Genome of the Chinese tree shrew. Nat Commun 4, 1426 (2013).

14. Fan, Y., et al. Chromosomal level assembly and population sequencing of the Chinese tree shrew genome. Zool Res 40, 506–521 (2019).

15. Xu, L., Chen, S.Y., Nie, W.H., Jiang, X.L. & Yao, Y.G. Evaluating the phylogenetic position of Chinese tree shrew (Tupaia belangeri chinensis) based on complete mitochondrial genome: implication for using tree shrew as an alternative experimental animal to primates in biomedical research. J Genet Genomics 39, 131–137 (2012).

16. Cao, J., Yang, E.B., Su, J.J., Li, Y. & Chow, P. The tree shrews: adjuncts and alternatives to primates as models for biomedical research. Journal of medical primatology 32, 123–130 (2003).

17. Park, U.S., et al. Mutations in the p53 tumor suppressor gene in tree shrew hepatocellular carcinoma associated with hepatitis B virus infection and intake of aflatoxin B1. Gene 251, 73–80 (2000).

18. Li, R., et al. Tree shrew as a new animal model to study the pathogenesis of avian influenza (H9N2) virus infection. Emerg Microbes Infect 7, 166 (2018).

19. Sanada, T., et al. Avian H5N1 influenza virus infection causes severe pneumonia in the Northern tree shrew (Tupaia belangeri). Virology 529, 101–110 (2019).

20. Yang, Z.F., et al. The tree shrew provides a useful alternative model for the study of influenza H1N1 virus. Virol J 10, 111 (2013).

21. Zhang, L., et al. Infectivity of Zika virus on primary cells support tree shrew as animal model. Emerg Microbes Infect 8, 232–241 (2019).

22. Wu, A., et al. Genome Composition and Divergence of the Novel Coronavirus (2019-nCoV) Originating in China. Cell host & microbe 27, 325–328 (2020).

23. Zhao, J., et al. Relationship between the ABO Blood Group and the COVID-19 Susceptibility. medRxiv, 2020.2003.2011.20031096 (2020).

24. Canrong, W., et al. In Silico Analysis of Intermediate Hosts and Susceptible Animals of SARS-CoV-2. ChemRxiv., Preprint. https://doi.org/10.26434/chemrxiv.12057996.v12057991 (2020).

25. Yang, J., et al. Prevalence of comorbidities and its effects in coronavirus disease 2019 patients: A systematic review and meta-analysis. International journal of infectious diseases: IJID: official publication of the International Society for Infectious Diseases 94, 91–95 (2020).

26. Lu, X., et al. SARS-CoV-2 Infection in Children. New England Journal of Medicine 382, 1663–1665 (2020).

27. Xu, Y., et al. Characteristics of pediatric SARS-CoV-2 infection and potential evidence for persistent fecal viral shedding. Nat Med 26, 502–505 (2020).

28. Yang, Y., et al. Exuberant elevation of IP-10, MCP-3 and IL-1ra during SARS-CoV-2 infection is associated with disease severity and fatal outcome. medRxiv, 2020.2003.2002.20029975 (2020).

29. Zhang, H., Penninger, J.M., Li, Y., Zhong, N. & Slutsky, A.S. Angiotensin-converting enzyme 2 (ACE2) as a SARS-CoV-2 receptor: molecular mechanisms and potential therapeutic target. Intensive Care Med Med 2020 04;46(4).

30. Damas, J., et al. Broad Host Range of SARS-CoV-2 Predicted by Comparative and Structural Analysis of ACE2 in Vertebrates. bioRxiv, 2020.2004.2016.045302 (2020).

31. Chen, L., Li, X., Chen, M., Feng, Y. & Xiong, C. The ACE2 expression in human heart indicates new potential mechanism of heart injury among patients infected with SARS-CoV-2. Cardiovasc Res (2020).

